# Integrated genomic and epigenetic profiling reveals RAS pathway as a key driver of odontogenic tumors

**DOI:** 10.1101/2025.09.09.675228

**Authors:** Miguel Ruiz-De La Cruz, Clara Estela Díaz-Velásquez, Norma Gabriela Resendiz-Flores, Héctor Martínez-Gregorio, Sebastian Galeana García, José Francisco Gómez Clavel, Cynthia Georgina Trejo Iriarte, Carlos Licéaga Escalera, Luis Ignacio Terrazas, Alejandro García Muñoz, Felipe Vaca-Paniagua

**Author notes:** Correspondence: Felipe Vaca Paniagua, Alejandro García Muñoz. These authors contributed equally (Miguel Ruiz-De La Cruz, Clara Estela Díaz-Velásquez).

## Abstract

Odontogenic tumors (OTs) are highly proliferative lesions with a low number of driving mutations, suggesting that concurrent alternative molecular mechanisms could support their extensive proliferation capacity. In this study, we analyzed 94 tissue samples from 79 patients with OTs and 15 healthy controls to explore their genetic and epigenetic alterations. Whole Exome Sequencing identified the *BRAF* V600E mutation in 75% of patients. A mutational hotspot analysis of six key genes (*NRAS*, *EGFR*, *BRAF V600E*, *SMO* L412, *SMO* W535, *KRAS* Q22K, *PTCH1* R602*, and *PTCH1* W129) revealed a high mutational rate, particularly in *BRAF* (91%), with 90% of *BRAF* V600E-positive patients being ameloblastomas. DNA methylation in the promoters of nine tumor-related genes (*RB1*, *RASSF1A*, *BRCA1*, *BRCA2*, *MSH2*, *MLH1*, *MGMT*, *TIMP3*, *BRAF*) was assessed in 67 OT patients and 15 controls. Five CpG sites showed significant hypermethylation (p<0.05; FDR q<0.05), notably in *RASSF1A* (cg50378469, cg50378539) and *TIMP3* (cg33197381, cg33197394, cg33197400). Somatic hypermethylation of the full promoter was detected in *RASSF1A* (8 patients, mean methylation: 17.5%), *BRCA1* (3 patients, mean methylation: 11.7%), and *MLH1* (1 patient, mean methylation: 2%). Interestingly, 76.5% of *BRAF* V600E-positive patients had *RASSF1A* promoter hypermethylation. A strong correlation between *BRAF* V600E mutation and *RASSF1A* hypermethylation was detected. Our results might imply a synergistic effect of the *BRAF* V600E mutation and *RASSF1A* hypermethylation as determinants in the RAS pathway. These findings, together with observations from other studies, suggest that the RAS pathway is a key axis in OT biology.

## INTRODUCTION

Odontogenic tumors (OTs) are rare, complex, and highly heterogeneous neoplasms originating from benign lesions of dental tissue, predominantly within bone, leading to the development of tumor masses that severely impact the quality of life of patients [1]. The World Health Organization (WHO) proposed a classification system in 2017 for odontogenic and maxillofacial bone tumors, based on their benign or malignant behavior and their epithelial, mesenchymal, or mixed origin [2], which are the most commonly used criteria for their characterization. OTs constitute approximately 5% of oral neoplasms, with about 95% being benign: 75% classified as ameloblastomas, odontomas, and myxomas, and close to 5% being malignant [3]. The precise etiology of OTs remains unclear. Nonetheless, some studies have employed next-generation sequencing to identify common specific mutations in OTs [4], [5], [6], while only a limited number have conducted Whole Exome Sequencing (WES) [7], [8].

The hallmark of OTs, extensive proliferation capacity, arises from the progressive accumulation of a handful of mutations that confer evolutionary adaptation to tumor cells. Altered signaling pathways in OTs include: i) Overactivation of the Ras pathway (Mitogen-Activated Protein Kinase pathway) driven by mutations in *BRAF* V600E and *KRAS* [9], [10], [11]; ii) ligand-independent activation of the β-Catenin pathway caused by mutations in *APC* and *CTNNB1* [12], [13], [14], [15]; iii) aberrant upregulation of the Hedgehog (Shh) pathway, resulting from mutations in *SMO* and *PTCH1*, as well as overexpression of its downstream effectors including as *GLI1* and *CCND1.* The Hedgehog pathway plays a key role in epithelial–mesenchymal interactions and cell proliferation during tooth development, and its dysregulation contributes to the progression of epithelial OTs [16], [17], [18], [19].

Although epigenetic mechanisms were initially overlooked, it is now known that they contribute significantly to the development of OTs due to their ability to alter gene expression independently of mutations in the coding region [20]. Nevertheless, research in this field has been limited. The molecular contribution of dysregulated epigenetic mechanisms involved in the formation and development processes of OTs includes three main groups: non-coding RNAs (ncRNAs) [21],[22],[23],[24],[25], histone post-translational modifications [26], and DNA methylation [27].

DNA methylation is the most studied epigenetic modification in OTs and frequently occurs in the promoter regions of tumor suppressor genes and oncogenes associated with the OTs development [28]. These aberrant methylation patterns in tumor tissue are widely recognized as somatic hypermethylation [29]. The evidence of these altered methylation marks can be classified into three main groups: i) hypomethylation in apoptosis-related genes (*TNFRSF25* and *BCL2L11*) [30], matrix metalloproteinases (*MMP-9*) [31], and cell cycle in odontogenic myxoma (*CDKN1B*, *TP53*, and *RB1*) [32], as well as hypomethylation of repetitive sequences such as long interspersed element-1 (*LINE-1*) [33]; ii) hypermethylation of tumor suppressor genes involved in cell cycleregulation, including *CDKN1A* in odontogenic keratocyst [34] and *CDKN2A* in ameloblastic carcinoma [35],[36]; and iii) overexpression of proteins involved in methyl group transfer to DNA (DNMTs), which further promotes aberrant methylation patterns [37].

Therefore, the aim of this study was to perform a comprehensive molecular analysis involving whole exome sequencing (WES), hotspot analysis, and methylation profiling of genes associated with OTs development. Specifically, we sought to identify novel aberrant methylation patterns at CpG sites in tumor tissue and explore their correlation with mutations in genes implicated in OTs pathogenesis. We employed advanced Next-Generation Sequencing (NGS) techniques targeting both coding regions (whole exome and hotspots) and non-coding regions (promoter methylation status), encompassing 79 patients and 15 healthy controls. Our findings highlight the critical role of the RAS pathway in the pathogenesis of these tumors.

## METHODS

### Study population

We selected 79 Mexican patients with fresh samples of odontogenic epithelial tumors, obtained via incisional biopsy, enucleation, or resection of the lesion, and 15 healthy population controls with samples collected from healthy oral tissues (gum, pulp, tooth, germ, or surrounding tissue) at the Almaraz Dental Clinic in Cuautitlán, Mexico, during the period from 2009 to 2019. All samples were immediately stored at -70°C for subsequent analysis. The inclusion criteria for the study were as follows: i) Availability of more than 2 µg of genomic DNA (gDNA). For the methylation profiling analysis, 12 patient samples were excluded due to low yield, ii) Samples obtained without prior chemotherapy or other pharmacological treatments.

### Ethics Statement and Patient Consent

All participants provided written informed consent for the use of their biological samples for research purposes. The study was conducted in accordance with the Declaration of Helsinki and was approved by the Ethics and Biosafety Committee of the Juárez Hospital of Mexico, Department of Maxillofacial Surgery, under protocol number HJM0623/19-1 (**Figure 1**).

**Figure 1.**
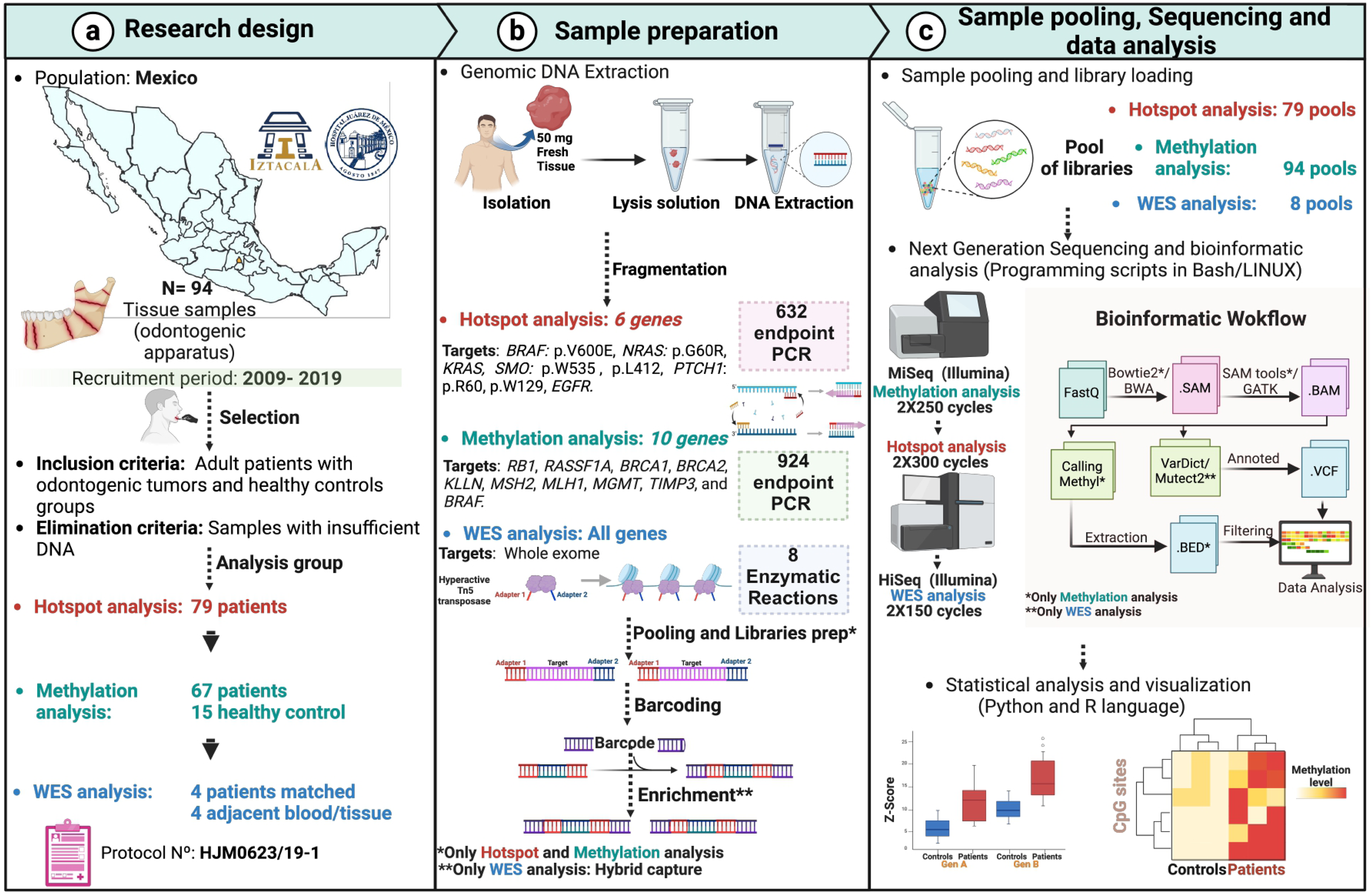
Experimental strategy. The recruitment and inclusion criteria for patient selection are shown in (**a)**. Sample preparation and treatment, the targeted for methylation, hotspot and whole exome Sequencing analysis, and total amplicons analyzed are presented in (**b**). The DNA library preparation workflow, barcoding, bioinformatics analysis, and biostatistical analyzes are represented in (**c**). Created with BioRender.com.

### gDNA extraction from tissue

gDNA was extracted from 50 mg of fresh tissue or 100 µL of peripheral blood using the DNeasy Blood & Tissues kit (Qiagen, Hilden, Germany). The integrity of the gDNA was assessed by electrophoresis on a 0.8% agarose gel, and purity was measured with an EPOCH BIOTEK spectrophotometer. Quantification was performed using a Quantus Fluorometer with the dsDNA Quantifluor kit (Promega, Madison, USA) (**Figure 1**).

### Positive methylation controls

Two positive controls for methylation were utilized: i) METC1-POS, a commercial Human HCT116 DKO Methylated DNA (Zymo Research, Irvine, CA, USA); and ii) METC2-POS gDNA, both of which were treated in vitro with the DNA methyltransferase enzyme M. *SssI*. These controls showed methylation percentages ranging from 98-100%, indicating complete methylation. The methylation status of these positive controls was confirmed using a methylation-sensitive restriction enzyme assay with *HpaII* (New England Biolabs, UK). Methylated CpG sites prevent the activity of *HpaII*. The positive controls have been previously validated and applied in our group [38].

### Methylation assay by bisulfite conversion

1000 ng of gDNA from 67 patients and 15 controls were processed using the EZ DNA Methylation-Gold kit (Zymo, California, USA) for sodium bisulfite conversion. The converted DNA was then quantified using the ssDNA Quantifluor kit (Promega, Madison, USA) and verified with a nanophotometer spectrophotometer (Implen). The bisulfite-treated DNA was subsequently stored at −20 °C, ready for targeted bisulfite sequencing.

### Primer design for bisulfite sequencing PCR and hotspot gene analysis

We designed eight pairs of primers targeting hotspot mutations in six genes involved in OT development: *NRAS*, *EGFR*, *BRAF V600E*, *SMO* L412, *SMO* W535, *KRAS*, *PTCH1* R602, and *PTCH1* W129. Primer design was conducted using Primer3 (v0.40) and specificity was verified using Primer-BLAST. The primers were further validated *in silico* using UCSC PCR tools (**Supplementary Table 2).**

Additionally, we designed primers for site-specific methylation analysis of the promoter regions of ten genes: *RB1*, *RASSF1A*, *BRCA1*, *BRCA2*, *KLLN*, *MSH2*, *MLH1*, *MGMT*, *TIMP3*, and *BRAF*. This was accomplished using MethPrimer 2.0 and Zymo Research online tools. For detailed primer design criteria, refer to the **Supplementary Table 1**.

### Endpoint PCR amplification

We performed a total of 924 endpoint PCRs to amplify the target regions within the promoters of the 10 genes (one amplicon per promoter) for the methylation analysis; and 632 endpoint PCRs to amplify hotspots regions in 6 genes. GoTaq Polymerase Master Mix® (Promega, Madison USA) was used for the amplification. Each reaction was performed in 25 µL as follows: 12.5 µL of GoTaq 2X Mix enzyme, primer forward and reverse were added to 200 nM, 20 ng of DNA converted with sodium bisulfite template for bisulfite direct PCR (methylation analysis) or 10 ng of non-converted DNA (hotspot analysis) and adjust to 25 µL total with RNAse-free water. A no-template reaction was used as a negative control. *MLH1* promoter region was used a positive control for methylation analysis.

The thermal cycling conditions used for the PCR were: one initial denaturation cycle at 95°C for 2 minutes; 34 cycles of denaturation at 95°C for 30 seconds, alignment at the primer specific Tm for 30 seconds, extension at 72°C for 30 seconds; a final extension step at 72°C for 5 min, and the reaction was kept at 4°C. Subsequently, the amplified products were resolved on a 0.8% agarose gel (**Supplementary Table 1-2)**.

### Pooling, library preparation and Next Generation Sequencing

Densitometric analysis was performed for each PCR product using the ImageLab® software and the amplicons were quantified by Fluorometry, equalized at 10 nM (lower concentration limit), and pooled by patient. We obtained 173 equimolar equalized PCR pools (79 pools for hotspot analysis and 94 pools for methylation analysis) with 6 genes for hotspot analysis and 11 genes for methylation analysis (9 study genes and 2 internal control genes). Each pool was purified with AMPure XP Beads (1.8X) and a total of 50 ng of DNA was used for the preparation of DNA libraries using the NEBNext® Ultra™ II DNA Library Prep Kit for Illumina®. The preparation of DNA libraries for WES analysis were done via hybrid capture enrichment using the Nextera Rapid Capture kit, SureSelectOxt and Exome IDT. Unequivocal molecular labeling for each of the samples was performed using NEBNext® Multiplex Oligos for Illumina® (index primers P5 and P7) using reduced (10X) cyclic amplifications. The generation of high-quality DNA libraries was confirmed by Bioanalyzer 2100 High Sensitivity DNA ChiP analysis, (Agilent, California USA). The samples were sequenced on two platforms for hotspot analysis with an experimental depth of 62X (**Supplementary** Figure 2a) using 2X300 cycles, and methylation analysis with an experimental depth of 1000X (**Supplementary** Figure 2b) using 2X250 cycles, both on a Miseq Illumina equipment. HiSeq Illumina equipment was used for WES analysis with 2x150 cycles.

### Bioinformatics analysis

Sequencing reads were processed through a bioinformatics pipeline, involving alignment using Bowtie2/BWA, SAM/BAM file generation with SAMtools/GATK (Bowtie2 and SAMtools were used for methylation analysis), and variant calling via BISMARK (for methylation analysis) or Mutect2/HaplotypeCaller (for hotspot and WES analysis), the variants were annotated with Annovar. Bases with Phred quality score >30 were kept for downstream analysis. Mapping and calling methylation were done with Bismark v0.22.3, using the GRCh37/hg19 genome reference. Methylation analysis were performed with a fully automatized workflow previously designed by our group [38]. Methylation analysis post-processing included annotation of variants, extraction of relevant data, filtering, and visualization.

### Statistical analysis

Statistical analysis and visualization were performed using Python v3.10.6 and R v4.1.2. The output methylation values were expressed as percentages (0-100%), extracted from a BED target file and normalized using the control samples by applying a Z-score for each specific CpG site. This normalization strategy has been previously employed in cancer studies to assess the methylation status of CpG sites in case-control studies. Patients were considered hypermethylated when their mean methylation (normalized z-score) exceeded the one-sided 95% confidence interval (CI) by 0.05. To control the Type I error rate in multiple comparisons, we performed the Wilcoxon test with Bonferroni correction on the p-values of all CpG sites. This analysis was carried out using scipy v1.10.1 and statsmodels v0.14. Differences with a p-value of less than 0.05 were considered statistically significant. Additionally, to correct for multiple testing, the false discovery rate (FDR) threshold was set at q<0.05 using the qvalue v2.32.0 library. A data matrix was constructed with pandas v1.5.2, and graphical representations were generated using ComplexHeatmap v2.13.1 (for methylation, WES, and hotspot data). The boxplots for methylation levels were created with ggplot2 v3.4.2.

## RESULTS

### 1. Clinical, epidemiological, and histopathological features of the study

The study includes 94 tissue samples from the odontogenic apparatus: 79 samples from patients with odontogenic epithelial and mesenchymal lesions, who underwent incisional biopsy to confirm the presumptive diagnosis, followed by surgical intervention for definitive diagnosis. The sample distribution was nearly equal, with a 1:1 ratio of 37 males (46.8%) and 38 females (48.1%), with 4 (5.1%) remaining unidentified (**Table 1**). The patients underwent enucleation of the lesion and curettage under general anesthesia (**Supplementary** Figure 1). Additionally, we included 15 healthy population controls with samples collected from healthy oral tissues (**Figure 1**).

**Table 1.** Epidemiological features and tissue sample types of the participants.

| Epidemiological features |  | Patients |  | Controls |  |
| --- | --- | --- | --- | --- | --- |
|  |  | N= 79 | 100% | N=15 | 100% |
| Age | | Mean ± S.D. | n (%) | Mean ± S.D. | n (%) |
| | <20 years | 15.7 ± 2.1 | 26 (32.9) | N/A | 1 (6.7) |
| | 20-40 years | 28.3 ± 6.1 | 25 (31.7) | 23.4 ± 2.3 | 11 (73.3) |
| | >40 years | 54.4 ± 11.0 | 20 (25.3) | 45 ± 4 | 3 (20) |
|  | Unknown | N/A | 8 (10.1) | N/A | 0 (0) |
| Sex |  | n | % | n | % |
|  | Male | 37 | 46.8 | 5 | 33.3 |
|  | Female | 38 | 48.1 | 10 | 66.7 |
|  | Unknown | 4 | 5.1 | 0 | 0.0 |
| Tissue type |  | Patients |  |  |  |
|  |  | N=79 | 100% |  |  |
| Lesion |  | n | % |  |  |
|  | **Ameloblastoma** | **23** | **29.1** |  |  |
|  | Solid | 11 | 13.9 |  |  |
|  | Unicystic | 11 | 13.9 |  |  |
|  | Plexiform | 1 | 1.3 |  |  |
|  | **Myxoma** | **10** | **12.7** |  |  |
|  | **Cyst** | **29** | **36.7** |  |  |
|  | Odontogenic keratocyst | 16 | 20.3 |  |  |
|  | Dentigerous | 5 | 6.3 |  |  |
|  | Residual | 1 | 1.3 |  |  |
|  | Odontogenic | 2 | 2.5 |  |  |
|  | Orthokeratinised odontogenic | 1 | 1.3 |  |  |
|  | Keratocystic odontogenic tumor | 4 | 5 |  |  |
|  | **Granuloma** | **2** | **2.5** |  |  |
|  | **Fibroma** | **5** | **6.3** |  |  |
|  | Ameloblastic fibroma | 1 | 1.3 |  |  |
|  | Ossifying fibroma | 3 | 3.7 |  |  |
|  | Myofibroma | 1 | 1.3 |  |  |
|  | **Adenomatoid odontogenic tumor** | **2** | **2.5** |  |  |
|  | **Other benign lesions<sup>a</sup>** | **8** | **10.2** |  |  |
| Tissue type |  | Controls |  |  |  |
|  |  | N=15 | 100% |  |  |
| Normal |  | n | % |  |  |
|  | Gum | 7 | 46.7 |  |  |
|  | Tooth | 6 | 40 |  |  |
|  | Germ | 1 | 6.6 |  |  |
|  | Surrounding | 1 | 6.6 |  |  |

<sup>a</sup>These samples of benign lesions include: 2 Benign nerve sheath tumors, 2 Benign osteoblastoma, 2 Lichen planus cases, 1 Intraosseous schwannoma, and 1 Submaxillary gland lesion.

There was a significant difference in the average age of patients and controls (31.3±17.3 vs. 27.5±10.6; p=0.042, t-test). The clinicopathological characteristics among patients included odontogenic epithelial tumors such as 23 ameloblastoma (29.1%), 29 cystic tumors (36.7%), 10 myxomas (12.7%), 5 fibromas (6.7%), 2 granulomas (2.5%), 2 adenomatoid OTs (2.5%), and 8 other benign lesions (10.2%). The healthy control tissue samples included 7 gum (46.7%), 6 tooth (40%), 1 germ (6.6%), and 1 surrounding (6.6%) (**Table 1**).

### 2. Hotspot and Whole Exome Sequencing analysis confirm common *BRAF* V600E mutation in odontogenic tumors

Each odontogenic lesion was classified according to international criteria (WHO, 2017) during the histopathological examination of the tumors (**Figure 2a-d, Supplementary** Figure 1). To evaluate the molecular pathogenesis underlying OT, we performed WES analysis on four patients paired with four adjacent tissue or blood controls. We identified the highly reported *BRAF* V600E mutation in three patients (75%), while one patient (25%) had the *BRAF* V600E mutation along with alterations in *FAM53A*, *ZFR*, *SACS*, *TANGO6*, and *L3MBTL4*. Additionally, in a patient without *BRAF* V600E mutation, we observed low-frequency alterations (25%) in *VLDRL* and *AGPS* (**Figure 2e**).

**Figure 2.**
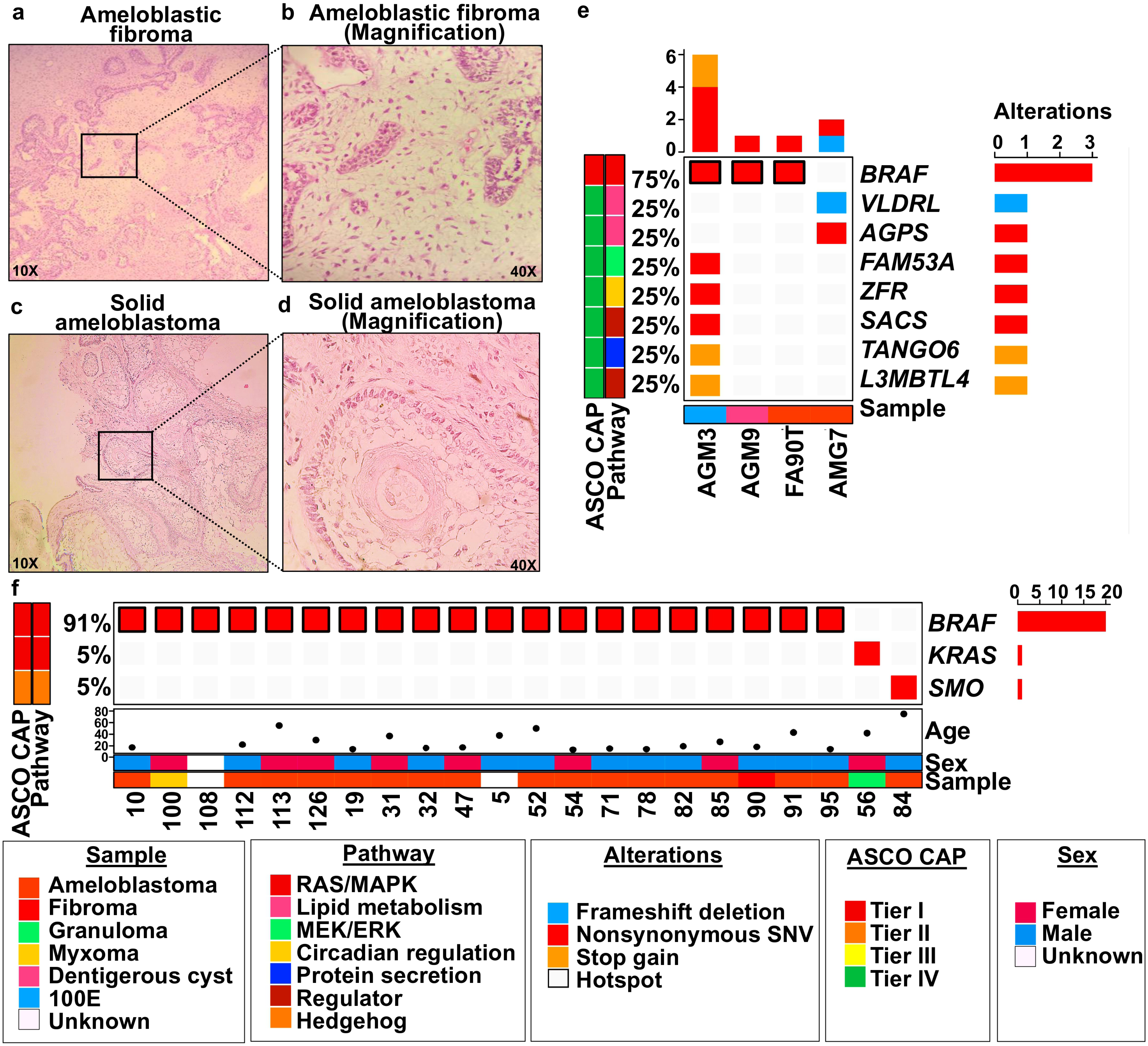
Mutational profiling of hotspot genes and Whole Exome Sequencing identifies *BRAF* V600E mutations as primary driver in odontogenic tumors. Two representative images of OTs are shown: **(a)** ameloblastic fibroma with rounded islands and narrow cords of odontogenic epithelium in a cellular mesenchymal background. **(b)** Ameloblastic fibroma at higher magnification (40X), reveals strands and islands of odontogenic epithelium with peripheral palisading nuclei resembling ameloblast-like cells and loosely arranged central cells similar to stellate reticulum, embedded in a myxoid, cell- rich stroma resembling dental papilla. **(c, d)** Histological patterns of solid ameloblastoma, specifically the follicular type, where angular cells often undergo cystic changes. This pattern resembles the epithelial component of the enamel organ within a fibrous stroma; peripheral cells are columnar to cuboidal with hyperchromatic nuclei arranged in a palisading pattern and reverse polarity, while central cells resemble stellate reticulum. **(e,f)** Mutations in *BRAF* V600E were predominant, found at high frequencies of 75% and 91% in our exome and hotspot analyses, respectively. Mutations in *KRAS* and *SMO* occurred at lower frequencies of 5%.

To better understand the genetic alterations implicated in OT development, we conducted a mutational hotspot analysis in a cohort of 79 patients, specifically targeting six genes that have been associated with these tumors: *NRAS*, *EGFR*, *BRAF V600E*, *SMO* L412, *SMO* W535, *KRAS* Q22K, *PTCH1* R602*, and *PTCH1* W129*. These mutations result in ligand-independent activation or loss of inhibition both in the RAS and Hedgehog pathways, which are critical in for the tumorigenesis process, and several studies have shown a significant synergistic crosstalk between these two pathways in malignant lesions [39].

Our results revealed that 22 out of the 79 patients (27.9%) harbored mutations in one or more of the targeted genes. Among the 22 mutations detected, 20 (91%) were *BRAF* V600E. Moreover, 18 of the 20 *BRAF* V600E mutations (90%) were found in patients with ameloblastomas. Additionally, mutations were detected in two other genes: *SMO* L412F (1/22, 4.5%) and *KRAS* Q22K (1/22, 4.5%) in individual cases, contributing to the mutational spectrum observed in this cohort (**Figure 2f, Supplementary Table 4**).

To refine the prevalence estimates of *BRAF* V600E mutations specifically in ameloblastomas, we focused on the 23 patients included in this study. Among these, 78.3% (18/23) presented the *BRAF* V600E mutation, reinforcing its significance as a driver mutation in this tumor type. Interestingly, one patient (ID FA90T/90), who was included in both the hotspot analysis and WES, consistently demonstrated the *BRAF* V600E mutation across both methodologies, highlighting the robustness and reliability of our findings.

### 3. Methylation profiling reveals crucial epigenetic marks in odontogenic tumor development

We assessed the methylation levels of the promoter regions of nine genes: eight tumor suppressor genes (TSGs) including *RB1*, *RASSF1A*, *BRCA1*, *BRCA2*, *MSH2*, *MLH1*, *MGMT*, *TIMP3*, and one oncogene, *BRAF*, in 67 patients with OTs and 15 healthy controls. The healthy controls were included to establish baseline methylation levels (**Figure 3, Supplementary Table 5**). Bisulfite sequencing PCR-NGS was used for this analysis. Additionally, *MLH1* and *KLLN* served as internal controls for bisulfite conversion and full methylation, respectively, as previously described by our group [38].

**Figure 3.**
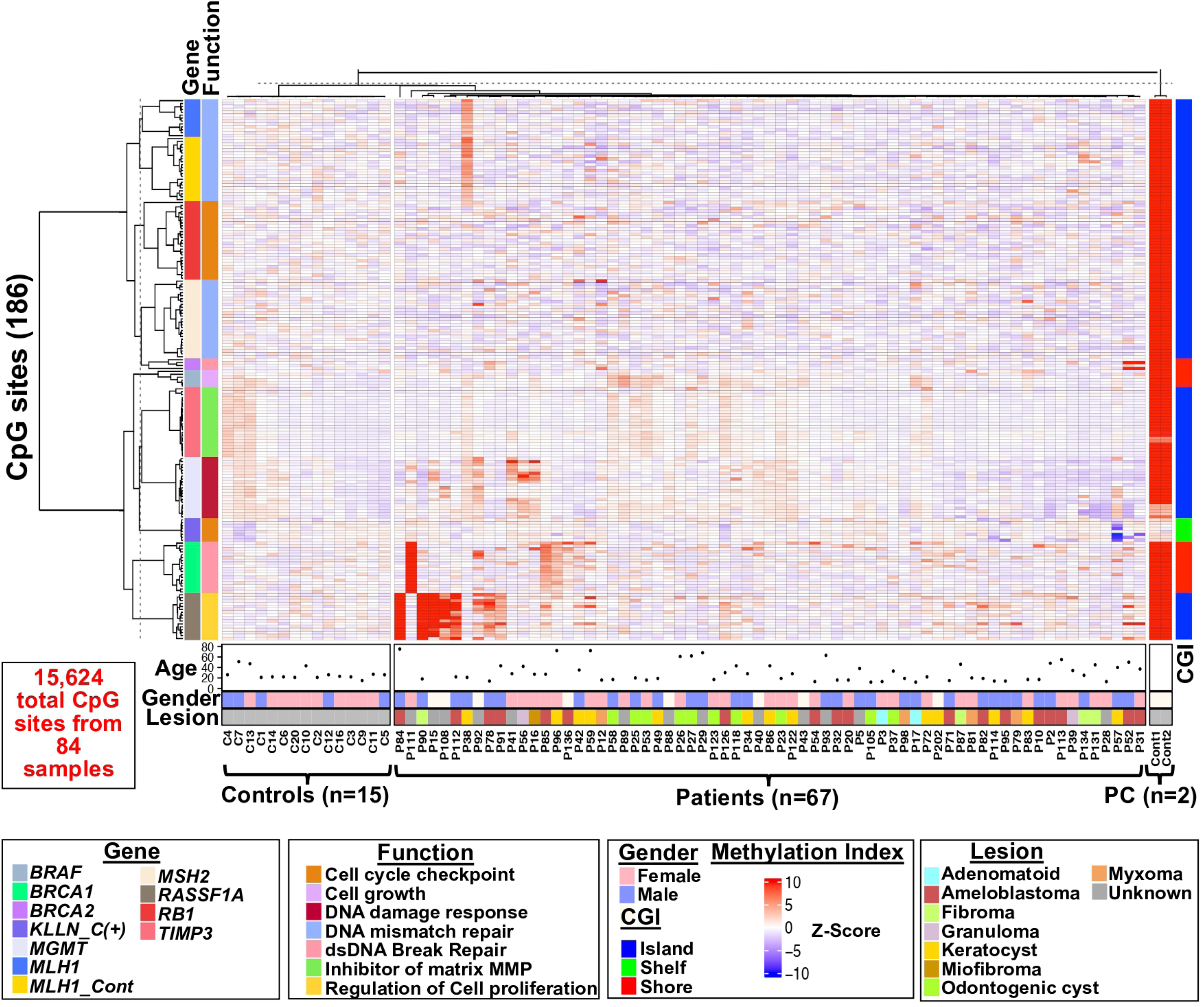
Methylation profiling reveals key epigenetic marks in odontogenic tumor. Site-specific methylation status in the promoter regions of the analyzed TSGs. The supervised hierarchical clustering shows 15,624 CpG sites from 186 individual CpG sites in tested genes (Y-axis) relative to case-control samples (X-axis). Gene function and CpG island distribution are illustrated in the left and right panels, respectively. Methylation values were normalized with the Z-score algorithm.

We mapped 186 CpG sites across nine locus-specific promoter genes (**Supplementary Table 3)**, resulting in a total of 15,624 CpG sites analyzed across all samples. A systematic evaluation using Revelio software revealed that none of the assessed CpG sites contained single nucleotide polymorphisms (SNPs) as analytical confounders (**Supplementary** Figure 3-4**; see Materials and Methods**). Furthermore, no SNPs in the target region were detected via the dbSNP track in the UCSC Genome Browser. We then calculated the average methylation values for all CpG sites (**Supplementary Table 5**) and identified five specific hypermethylated CpG sites (FDR q<0.05): three in *TIMP3* (**cg33197381**: Z-score 0.430 vs. -6.66E-11, *p*=0.03; **cg33197394**: Z-score 0.302 vs. 0, *p*=0.03; **cg33197400**: Z-score 0.148 vs. 6.66E-11, *p*=0.029) and two in *RASSF1A* (**cg50378469**: Z-score 4.376 vs. -6.66E-11, *p*=0.03; **cg50378539**: Z-score 3.675 vs. 6.66E-11, *p*=0.005) (**Supplementary Table 6**).

To further investigate the methylation patterns of the studied gene promoters and their correlation with patient data, we conducted a Pearson correlation (PC) analysis, which was projected onto a hierarchical clustering heatmap. By using the the normalized Z-scores of all CpG sites, the methylation profiles of gene promoters in both patients and controls revealed the segregation of two distinct patient clusters (**Figure 4a**). These clusters exhibited high methylation levels in the *RASSF1A* (6 patients) and *BRCA1* (4 patients) genes, both with a positive PC >0.6. The variance magnitude in the x-axis was 65.5, and 52.9 in the y-axis **(Figure 4a, Supplementary** Figure 5). This result was further confirmed through principal component analysis (PCA), comparing patient and control genes, where a clear separation was observed into two groups with the highest variation in methylation levels **(Figure 4b)**. These groups corresponded to 6 specific CpG sites in *RASSF1A* (cg50378539, cg50378469, cg50378445, cg50378413, cg50378515, and cg50378529) and *BRCA1* (cg41277462 and cg41277487). Only the sites cg50378539 and cg50378469 were found to be hypermethylated (**Figure 4c-d**, p<0.05; Wilcoxon signed-rank test**; Supplementary Table 8)**.

**Figure 4.**
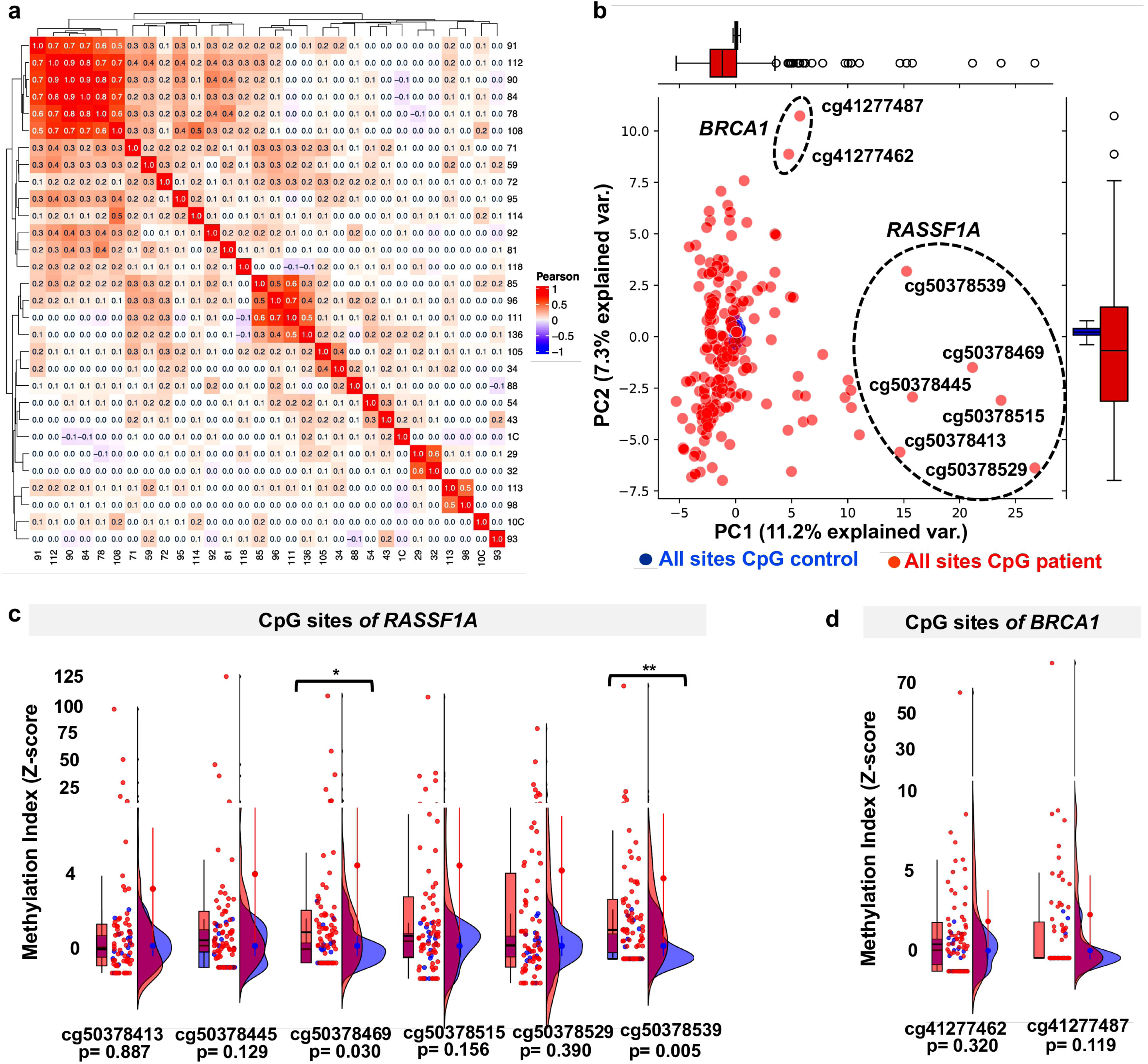
Methylation profiling identifies high methylation level in *RASSF1A* and *BRCA1* genes. **(a)** The unsupervised hierarchical clustering heatmap with Pearson correlation for each promoter (nine genes) reveals the separation of two patient groups associated with high methylation levels of the *RASSF1A* and *BRCA1* genes. **(b)** Principal component analysis of the mean Z-scores across 186 individual sites indicates that the variation in site-specific methylation levels is concentrated in two CpG sites of *BRCA1* and six of *RASSF1A*. **(c)** We identified two hypermethylated sites of *RASSF1A* in patients (cg50378469, p=0.03 and cg50378539, p=0.005; Wilcoxon signed-rank test), both statistically significant. **(d)** No significant differences in methylation were observed at the two *BRCA1* sites in the case-control analysis. Dotted lines indicate the k-means clustering for each group in **(b)**. One asterisk (*) denotes p <0.05, and a double asterisk (**) indicates p ≤0.005. Patients are shown in red and controls in blue in panels 4b-d.

### 4. Patients with odontogenic tumors showed locus-wide somatic hypermethylation in *RASSF1A*

A detailed analysis of locus-wide hypermethylation across the entire promoter revealed hypermethylation in three tumor suppressor genes: **i)** *RASSF1A*: 8 patients (mean methylation: 17.5%, range: 3.9–69.6%; 16 CpG sites); **ii)** *BRCA1*: 3 patients (mean methylation: 11.7%; range: 2.6–29%; 18 CpG sites); **iii)** *MLH1*: 1 patient with low methylation (mean methylation: 2%; 13 CpG sites) **(Figure 5a-b)**. Statistical analysis comparing patients vs controls for the *BRCA1* and *RASSF1A* promoters did not show significant differences **(Figure 5c-d)**.

**Figure 5.**
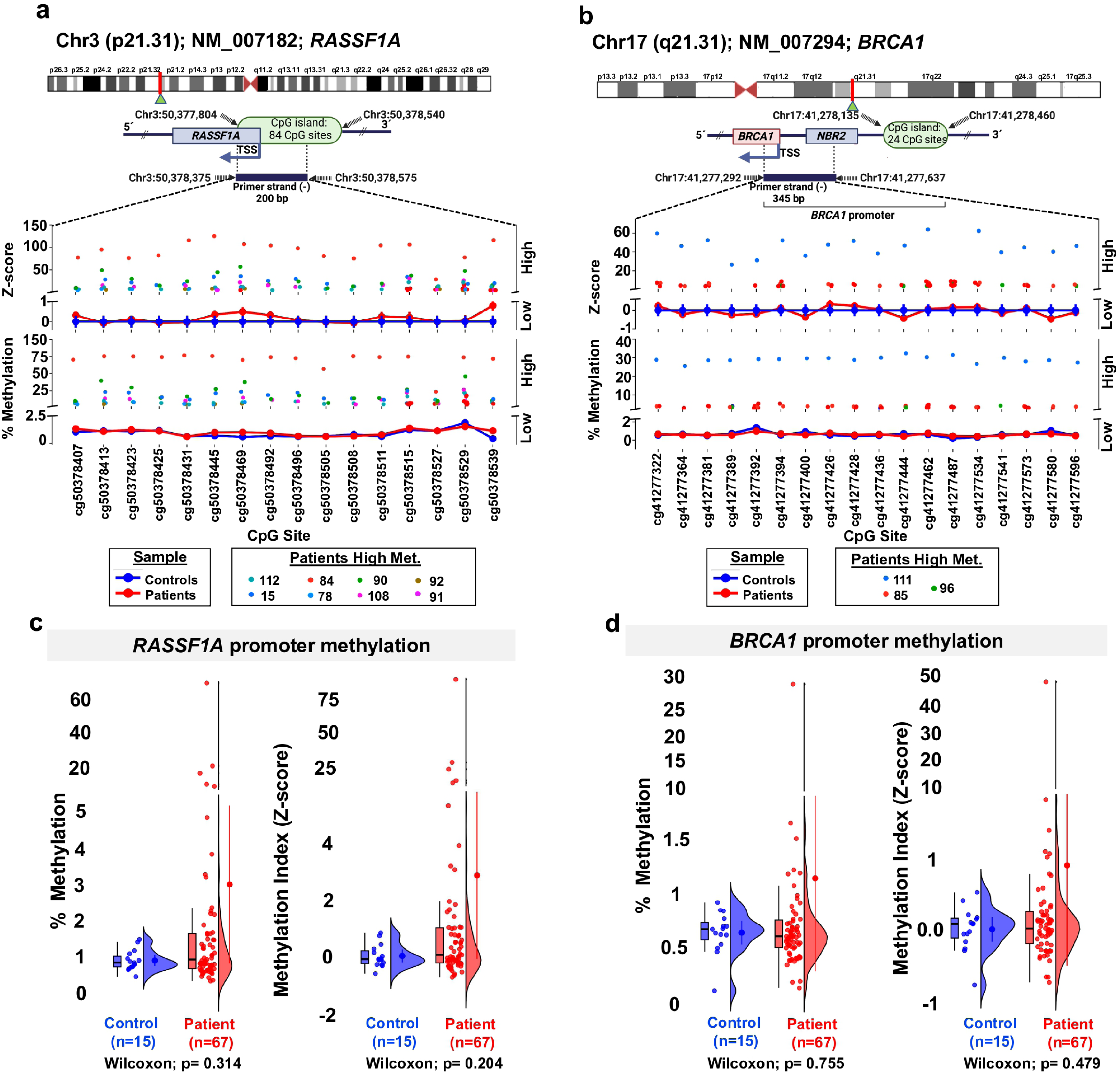
Comparison of site-specific somatic methylation levels across the promoter region of *RASSF1a* and *BRCA1* in patients vs controls. The methylation levels of the *RASSF1a* and *BRCA1* gene loci are shown in panels **(a)** and **(b)**, respectively. Somatic hypermethylation of the *RASSF1a* locus was observed in 8 patients, while hypermethylation of the *BRCA1* locus was observed in 3 patients. No significant differences were found between patient and control groups for these tumor suppressor genes (panels **c** and **d**, p>0.05). The solid blue line represents the average methylation values for DNA derived from blood controls, while the red line indicates the average methylation values for patients. Gene region schematics highlight the CpG island (CGI) and the transcriptional start site (TSS). All genes are shown in their polarity context as found in their *loci*.

Histopathologically, patients with hypermethylation across the *RASSF1A* locus included 4 ameloblastomas (50%), 2 fibromas (25%), and 2 other odontogenic lesions (25%). For *BRCA1*, the 3 female patients exhibited cystic lesions in the odontogenic apparatus **(Supplementary Table 7)**.

### 5. Mutational status of *BRAF* V600E in odontogenic tumors shows high correlation with *RASSF1A* promoter hypermethylation

To investigate the relationship between the *BRAF* V600E mutation and methylation status in the *RASSF1A* promoter, we found that of 76.5% (13/17) patients with the *BRAF* V600E mutation also had *RASSF1A* hypermethylation above 2.5% compared to the baseline level close to 0, with a strong correlation (PC >0.6). Among those with *RASSF1A* promoter hypermethylation, 85% (11/13) were diagnosed with ameloblastomas (**Figure 6a-b; Supplementary Table 7**). To further explore this connection, we compared the *BRAF* V600E mutational status with the *RASSF1A* promoter methylation levels in the patients. Notably, a significant difference was observed between the *BRAF* V600E mutation carriers, non-carriers, and controls (p<0.0001; Wilcoxon signed-rank test) (**Figure 6c; Supplementary** Figure 6).

**Figure 6.**
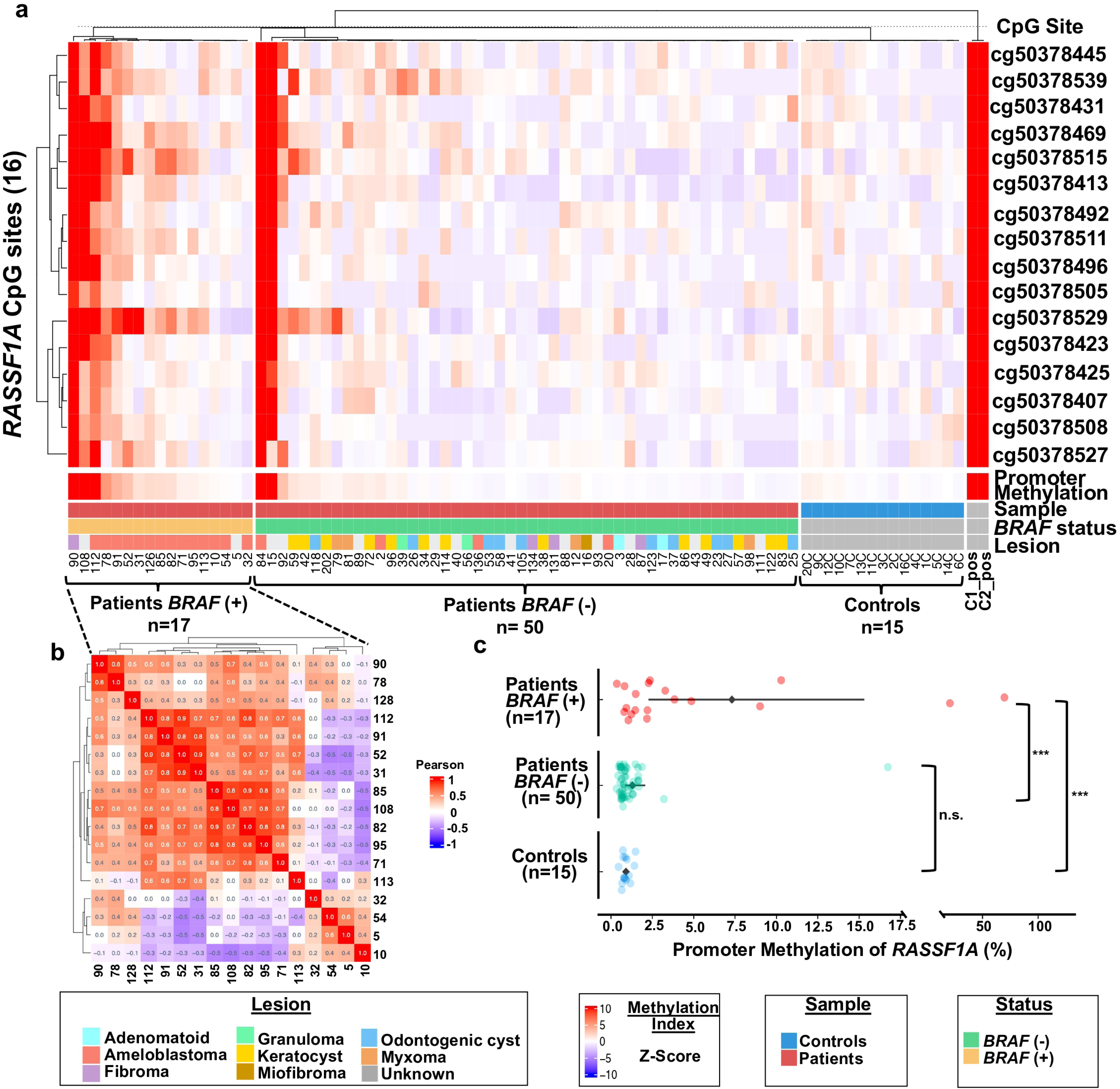
Correlation between oncogenic *BRAF* V600E status and *RASSF1A* promoter methylation. The supervised hierarchical clustering (based on the *BRAF* V600E mutational status) shows the site-specific methylation status in the promoter region of *RASSF1a* across 16 individual CpG sites in the tested genes (Y-axis) relative to case-control samples (X-axis) **(a)**. A high correlation between *BRAF* V600E mutational status and hypermethylation in *RASSF1a* was observed in 13 out of 17 patients **(b)**. Comparative analysis shown in panel **(c)** revealed statistically significant differences versus controls (p<0.0001). Methylation status, sample type, lesion, and *BRAF* status are illustrated in the bottom panels. Methylation values were normalized using the Z-score. *BRAF*(+): *BRAF V600E*.

## DISCUSION

OTs are rare lesions with an unclear etiopathogenesis and extensive proliferative capacity [40]. While most are of benign origin, they are characterized by local aggressiveness in the maxilla or mandible, high infiltrative potential, and a tendency for increased recurrence [41]. Ameloblastoma is the most common OT, originating from odontogenic epithelium [42]. It accounts for 19.3–41.5% of all OTs [43], [44]. In our study, we included 23 patients of ameloblastoma (29.1%) and 29 cystic tumors (36.7%).

Studies have focused on mutations in the *BRAF* gene, which is recognized as a critical activator of the MAP kinase (MAPK) pathway, driving cell cycle progression and cell growth [45]. The *BRAF* V600E mutation accounts to approximately 90% of all *BRAF* gene mutations and is implicated in 80-90% of ameloblastic lesions [46]. This detection is commonly performed using techniques such as immunohistochemistry [47], Sanger sequencing [48], or more recently, advanced methods like high-performance MALDI-TOF analysis [49]. In our study, we employed two powerful identification techniques: Hotspot and WES analysis using next-generation sequencing (NGS), confirming the prevalence of the common *BRAF* V600E mutation in OTs. Among the ameloblastoma cases, 78.3% exhibited the *BRAF* V600E mutation. *KRAS* mutations have been detected in a limited number of patients with adenomatoid OTs [9], [50], while L412F ligand-independent activation of *SMO* mutation has also been reported in ameloblastoma, occurring in 4-13% of patients [51],[52], or not detected [53]. We identified mutations in *KRAS* and *SMO* L412F in only 2.5% of the cases. Importantly, none of these mutations were in the germline. Notably, previous studies have reported significant crosstalk between the Hedgehog and RAS pathways, wherein GLI transcription factors, the final effectors of the Hedgehog pathway, are essential for RAS-induced proliferation in cancer [54]. Future studies should prioritize the functional exploration of the Hedgehog pathwaýs role in regulating RAS-induced proliferation in OTs.

Genetic and epigenetic changes may play a major role in the development of OTs, even though their exact molecular pathophysiology is unknown [20],[27]. Most prior studies used qualitative techniques, such as methylation-specific PCR, which lack the resolution and precision of newer methodologies [34]. To our best knowledge, this is the first study to evaluate DNA methylation levels in a broad sample of OTs, analyzing nine genes involved in tumorigenesis (*RB1*, *RASSF1A*, *BRCA1*, *BRCA2*, *MSH2*, *MLH1*, *MGMT*, *TIMP3*, *BRAF*) by quantitative assessment of methylation levels using NGS, providing more accurate measurements. The sensitivity of this approach allowed for the identification of five novel hypermethylated CpG sites: three in *TIMP3* (cg33197381, cg33197394, cg33197400) and two in *RASSF1A* (cg50378469, cg50378539). These markers may play a key role in the epigenetic silencing of OTs at the somatic level.

Hypermethylation in OTs is associated with the overexpression of *DNMT3A* and *DNMT3B* [37], [55] and has been reported in tumor suppressor genes in isolated cases, including *CDKN1A* (p21) in odontogenic keratocysts (3/10, 30% of patients vs. 0/10, 0% of dental follicles used as controls) [34], *CDKN2A* (p16) in ameloblastic carcinoma (single case report) [35], and in another study involving 8 patients (100%) vs. 6 controls (0% of dental follicles used as controls) [36]. Our study expands on this knowledge by identifying hypermethylation in three tumor suppressor genes not previously reported in OTs: *RASSF1A*, *BRCA1*, and *MLH1*. Hypermethylation of *RASSF1A* was found in 13/69 (19.4%) patients, with methylation levels ranging from 2% to 70%. Of these, 8 patients exhibited locus-wide hypermethylation (4-70% methylation level) across the 16 CpG sites evaluated. Importantly, 50% of ameloblastomas, 25% of fibromas, and 25% of other odontogenic lesions showed hypermethylation at this locus. The expression of *RASSF1A* is regulated by the recognition of three consensus GC boxes in the promoter region by Sp1 and Sp3 zinc finger domain transcription factors, with their DNA binding being inhibited by cytosine methylation [56], [57]. The loss of expression due to aberrant hypermethylation of *RASSF1A* is also commonly observed in the progression of multiple types of cancer and promotes Ras pathway upregulation [58]. Hypermethylation of *BRCA1* was detected in female patients with cystic lesions, including intraosseous cystic neuroma (29% methylation), unicystic ameloblastoma (3.5% methylation), and odontogenic keratocyst (2.6% methylation). Low levels of hypermethylation in *MLH1* (2% methylation) were observed in a single patient with odontogenic keratocyst.

Our study presents compelling evidence of a significant prevalence of *BRAF* V600E mutations and *RASSF1A* promoter hypermethylation in patients with odontogenic lesions. Notably, only one previous study has explored the association between these two genetic-epigenetic mechanisms, however it produced inconclusive results. The study qualitatively assessed *RASSF1A* methylation status and *BRAF* V600E immunohistochemical expression in 66 odontogenic lesions, finding no association. They reported *RASSF1A* hypermethylation in 20% (4/20) of cases, while 85.7% (18/21) of ameloblastomas were *BRAF* V600E-positive [59]. Conversely, our study used a more quantitative and sensitive NGS-based approach, analysing a larger cohort of patients, and found a significant correlation: 76.5% (13/17) of patients with the *BRAF* V600E mutation exhibited *RASSF1A* hypermethylation. We found that *RASSF1A* promoter hypermethylation seems to be a hallmark of ameloblastomas with a prevalence 85%. Previous reports have also found correlations between *RASSF1A* hypermethylation and *KRAS* mutations in colorectal cancer [60], as well as with *BRAF* mutations in thyroid cancer [61],[62]. Additionally, concomitant *RASSF1A* hypermethylation and *KRAS*/*BRAF* mutations have been reported to occur preferentially in MSI-high sporadic colorectal cancer, suggesting a potential synergistic effect [63].

RASSF1A interacts with Ras, potentially playing a role in mediating the tumor-suppressive effects within the Ras signaling pathway [64]. Our findings of *RASSF1A* promoter hypermethylation and its associated epigenetic silencing in odontogenic lesions suggest a potential functional disruption of this tumor-suppressive pathway. This silencing could lead to the loss of *RASSF1A* ability to regulate key downstream signaling events within the Ras cascade, thereby enhancing cell survival, proliferation, and resistance to apoptosis.

This study provides valuable insights on the genetic and epigenetic alterations in OTs; however, some limitations should be considered: i) The number of case and control samples was unequal. ii) The mutational status of *BRAF* V600E was not assessed in controls, though previous sequencing studies have not detected mutations in this gene in normal tissue [65]. iii) We did not evaluate *RASSF1A, BRCA1* and *MLH1* expression at the RNA or protein level, leaving open the question of functional silencing. This limitation is somewhat alleviated by prior landmark reports demonstrating loss of expression by epigenetic silencing of these genes in other tumors [58], [66], [67]. iv) We did not evaluate the role of *RASSF1A* neighboring gene *ZMYND10*, which is considered a proapoptotic and antiangiogenic tumor suppressor that is frequently hypermethylated and mutated in cancers [68].

We summarize our findings in three key events (**Figure 7**): i) genetic and epigenetic alterations reported in this study (**Figure 7a**), ii) the implications of oncogenic signaling in *BRAF* and its potential as a therapeutic target (**Figure 7b**), and iii) the impact of *RASSF1A* epigenetic inactivation and its potential synergism in the Ras-MAPK pathway (**Figure 7c**). Collectively, our findings indicate a key implication of the RAS pathway in OT.

**Figure 7.**
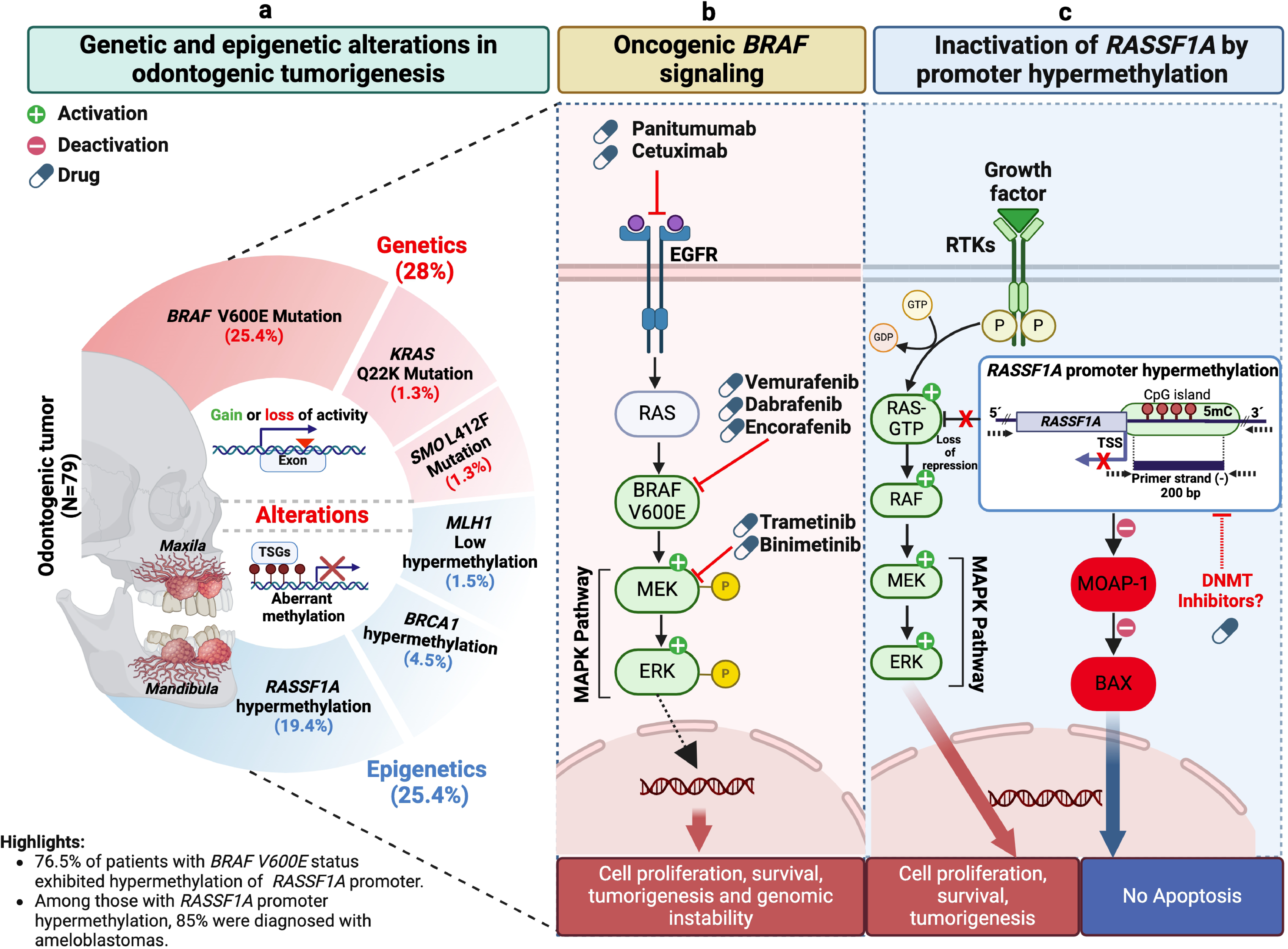
Integrated genomic and epigenetic model of RAS pathway as a key driver of odontogenic tumors. The summarized model of the genetic (*BRAF* V600E, *KRAS* Q22K, and *SMO* L412F) and epigenetic (promoter hypermethylation across the *RASSF1A*, *BRCA1*, and *MLH1 loci*) alterations identified in the study is shown in panel **(a)**. Oncogenic signaling of *BRAF* V600E **(b)** and promoter hypermethylation of the *RASSF1A* locus **(c)** are shared by approximately 80% of the patients, with the RAS-MAPK aberrant pathway, acting as the determinant in the etiopathogenesis of OTs. **5mC:** 5- Methyl Cytosine. **TSG:** Tumor Suppressor Gene. **EGFR:** Epidermal Growth Factor Receptor. **RTK:** Receptor Tyrosine Kinase. **DNMT:** DNA Methyltransferase. **TSS:** Transcription Start Site. Created with BioRender.com.

## CONCLUSION

Our results, together with findings from other studies, including (i) a significant mutational rate in *BRAF* and *RAS*, (ii) the concomitant methylation of *RASSF1A* and *BRAF* mutations, and (iii) Hedgehog pathway activation of RAS-induced proliferation, strongly support the hypothesis that the RAS pathway is the main driver of OTs. These findings highlight the RAS pathway as a central axis in OT biology, encouraging future studies to further define its contributions to tumor progression.

## DATA AVAILABILITY

The data presented in this study are available in the Sequence Read Archive repository under accession number PRJNA1211046 for the methylation analysis, PRJNA1207096 for the hotspot mutational analysis, and PRJNA1212530 for the whole exome sequencing (WES) analysis.

## CODE AVAILABILITY

The codes used in this study are deposited in https://github.com/UBIMED-Lab13/MethylationDetection.git.

## ACKNOWLEDGEMENTS

This work was supported by UNAM PAPIIT IN225224, UNAM PAPIIT IN225820, UNAM PAPIIT IN225920, CONACYT Fondo Sectorial 272573, Fondo SEP CONACYT 285879.

We would like to express our sincere gratitude to the Maxillofacial Surgery Department at Hospital Juárez de México for their valuable collaboration on this project.

## AUTHOR CONTRIBUTIONS

Conception, M.R.D.L.C., A.G.M. and F.V.P. Patient recruitment, sampling, database, A.G.M., M.R.D.L.C., S.G.G., L.B.N., J.F.G.C., C.G.T.I., C.L.E. Experimental analysis, M.R.D.L.C., C.E.D.V., N.G.R.F., S.G.G., L.B.N., and F.V.P. Data analysis and visualization, M.R.D.L.C., H.M.G., F.A.B., and F.V.P. Manuscript writing and review, M.R.D.L.C., H.M.G., C.E.D.V., F.A.B., N.G.R.F., A.G.M. and F.V.P. Resources, A.G.M. and F.V.P. Funding acquisition, F.V.P. All authors contributed to the article and approved the submitted version.

## COMPETING INTERESTS

The authors declare no competing interests.

